# Regulation of *Blos1* by IRE1 prevents the accumulation of Huntingtin protein aggregates

**DOI:** 10.1101/2021.09.28.462237

**Authors:** Donghwi Bae, Rachel Elizabeth Jones, Katherine M. Piscopo, Mitali Tyagi, Jason D. Shepherd, Julie Hollien

## Abstract

Huntington’s disease is characterized by accumulation of the aggregation-prone mutant Huntingtin (mHTT) protein. Here, we show that expression of mHTT in mouse cultured cells activates IRE1, the transmembrane sensor of stress in the endoplasmic reticulum, leading to degradation of the *Blos1* mRNA and repositioning of lysosomes and late endosomes toward the microtubule organizing center. Overriding *Blos1* degradation results in excessive accumulation of mHTT aggregates in both cultured cells and primary neurons. Although mHTT is degraded by macroautophagy when highly expressed, we show that prior to the formation of large aggregates, mHTT is degraded via an ESCRT-dependent, macroautophagy-independent pathway consistent with endosomal microautophagy. This pathway is enhanced by *Blos1* degradation and appears to protect cells from a toxic, less aggregated form of mHTT.

**Condensed title:** *Blos1* regulation protects from Huntingtin aggregation

**Short summary:** Here the authors demonstrate that the regulation of *Blos1* by the unfolded protein response prevents the excessive accumulation of Huntingtin, the protein that underlies Huntington’s disease. This pathway involves the macroautophagy-independent degradation of Huntingtin prior to its forming large aggregates.

## Introduction

Protein aggregation underlies several common and devasting neurodegenerative diseases, for which there are no cures (Arrasate and Finkbeiner, 2012; Gan *et al*., 2018). However, the distinctive protein deposits that form within the brain are not always correlated with the specific neurons whose death or loss of function leads to the symptoms experienced by patients (Tompkins and Hill, 1997; Gutekunst *et al*., 1999; Kuemmerle *et al*., 1999). This observation has led to the idea that smaller, oligomeric forms of aggregation-prone proteins are more likely to be toxic, by interfering with normal cellular processes (Ross and Poirier, 2005). In Huntington’s disease, for example, large protein aggregates of the mutant Huntingtin protein (mHTT) are thought to be somewhat protective (Saudou *et al*., 1998; Arrasate *et al*., 2004). These aggregates can be degraded by macroautophagy (MA), which encloses aggregates in double-membraned autophagosomes that then fuse with lysosomes for degradation (Iwata *et al*., 2005; Olzmann, Li and Chin, 2008). Whether smaller, potentially more toxic, oligomers are also degraded by MA or by distinct pathways is not clear.

Aggregating proteins tend to accumulate near the juxtanuclear microtubule organizing center (MTOC), as a result of minus end-directed trafficking on microtubules, a process that is dependent on the ubiquitin-binding adapter protein HDAC6 (Kawaguchi *et al*., 2003; Olzmann, Li and Chin, 2008). Disruption in trafficking hinders both the coalescence into larger aggregates and the degradation of these proteins (Kawaguchi *et al*., 2003). Minus-end directed trafficking of lysosomes has also been observed in cellular models of Huntington’s and Parkinson’s diseases (Dodson *et al*., 2012; Erie *et al*., 2015), although long-distance trafficking along axons is often disrupted in these diseases (Lie and Nixon, 2019).

We have found that clustering of lysosomes and late endosomes (LEs) near the MTOC can result from a specific pathway induced by stress in the endoplasmic reticulum (ER) (Bae *et al*., 2019). Inositol requiring enzyme 1 (IRE1), a conserved sensor of ER stress and key mediator of the unfolded protein response, is a nuclease that cleaves the mRNA encoding the transcription factor XBP1 to initiate its splicing and activation, leading to the upregulation of many genes involved in ER homeostasis. IRE1 also cleaves several other mRNAs, initiating their degradation through the Regulated IRE1-Dependent Decay (RIDD) pathway (Hollien *et al*., 2009; Moore and Hollien, 2015). One of the key targets of RIDD is the mRNA encoding biogenesis of lysosome-related organelles complex 1 subunit 1 (BLOC1S1, or BLOS1). BLOS1 is a member of the BLOC1-related complex (BORC), which links lysosomes to kinesin, allowing for their trafficking toward the periphery of the cell (Pu *et al*., 2015). BLOS1 is also important for several other trafficking and endosome-related functions, including the sorting of proteins in endosomal compartments (Zhang *et al*., 2014), kinesin switching during endosome recycling (Zhang *et al*., 2020), autolysosomal tubulation (Wu *et al*., 2021), and tubulin acetylation (Wu *et al*., 2018). Degradation of the *Blos1* mRNA by RIDD during ER stress leads to the repositioning of LE/ lysosomes to the MTOC and to the enhanced degradation of ubiquitinated proteins (Bae *et al*., 2019).

Because neurodegenerative diseases such as Huntington’s isease often display markers of ER stress (Hetz and Saxena, 2017; Shacham, Sharma and Lederkremer, 2019), we hypothesized that the degradation of *Blos1* may contribute to clearance of disease-associated aggregating proteins by trafficking degradative organelles to the cell center. Here we show that expression of the aggregation-prone Huntingtin protein induces degradation of *Blos1* mRNA, which in turn protects cells from apoptosis and prevents the accumulation of aggregates by enhancing an alternative pathway to degradation.

## Results & Discussion

To test whether expression of the mutant Huntingtin protein is affected by IRE1 and the RIDD pathway, we transfected mouse MC3T3-E1 cells with plasmids encoding exon 1 of either the wild-type Huntingtin protein (wtHTT), which contains a string of 23 Gln residues, or a disease-causing mutant Huntingtin (mHTT), which contains 145 Gln’s, each tagged with GFP. Cells expressing wtHTT-GFP displayed a diffuse, cytosolic expression pattern, whereas approximately 20% of cells expressing mHTT-GFP displayed large, juxtanuclear aggregates, as expected (Fig. 1A). These aggregates colocalized with the microtubule organizing center in 66% of the aggregate-containing cells (Fig. 1A). We then constructed stable MC3T3-E1 cell lines expressing the wtHTT-GFP or mHTT-GFP proteins under the control of a doxycycline-inducible promoter. We transfected HTT plasmids into cell lines expressing either *Rfp* (as a control) or a stabilized version of the *Blos1* mRNA (*Blos1*^*s*^), which contains a silent point mutation rendering it resistant to degradation by RIDD (Moore and Hollien, 2015). We previously showed that these *Blos1*^*s*^ - expressing cells override the repositioning of LE/lysosomes during ER stress (Bae *et al*., 2019). Addition of doxycycline (4.5 μM, 72 h) to control cells and the subsequent expression of mHTT, but not wtHTT, led to the activation of IRE1 as assessed by *Xbp1* splicing (Fig 1B), degradation of RIDD targets (Fig 1C, D), and repositioning of LE/lysosomes to the MTOC (Fig 1E, F). As expected, when we induced mHTT expression in cells expressing *Blos1*^*s*^, IRE1 was activated but LE/lysosomes did not reposition (Fig 1B-H).

**Figure 1.**
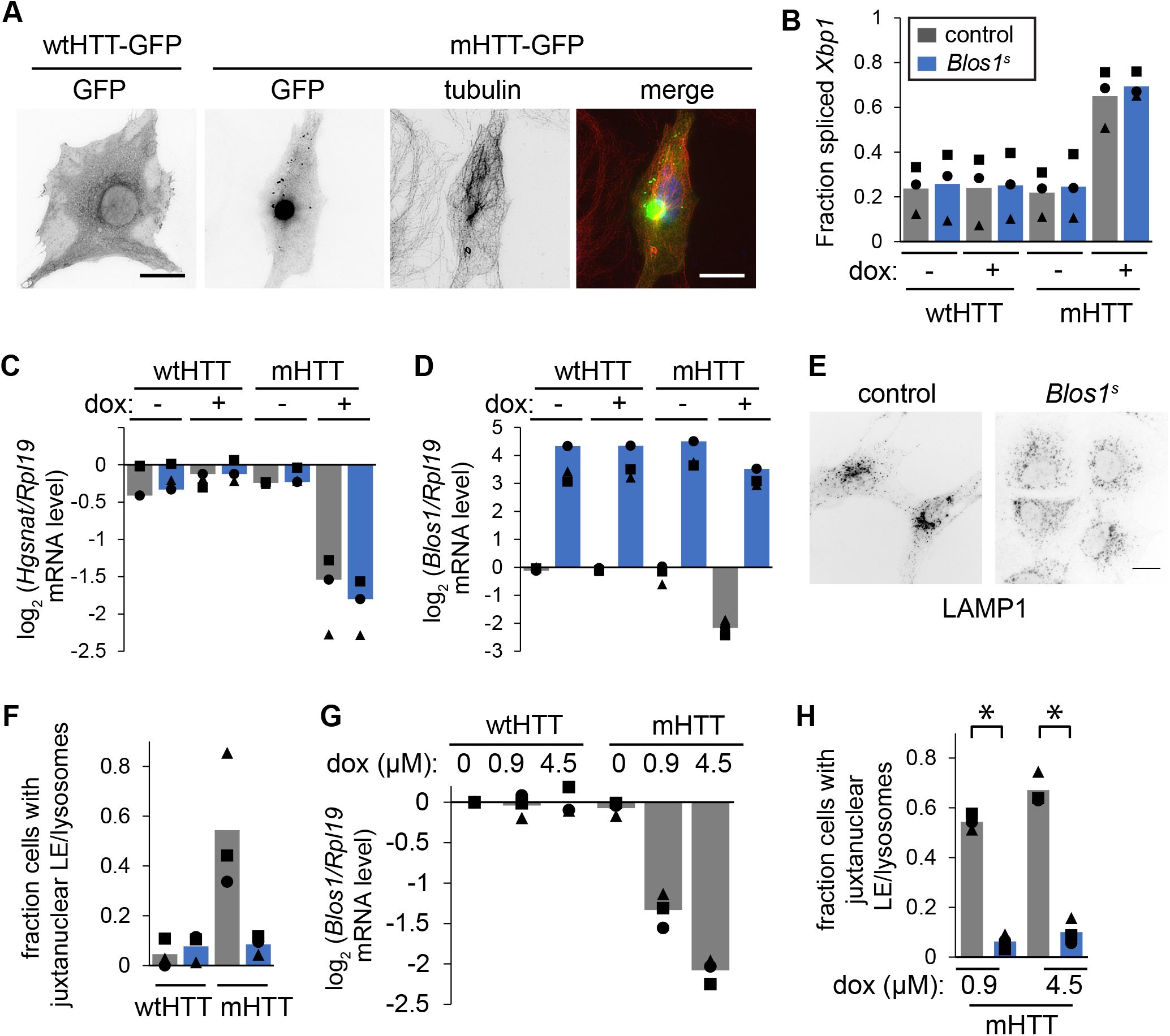
Expression of mHTT but not wtHTT induces IRE1. (A) We transfected MC3T3-E1 cells with either wt or mHTT-GFP, then fixed and stained using antibodies for GFP and alpha-tubulin (scale bar, 10 μm). (B) We constructed stable MC3T3-E1 cells expressing either *Rfp* (labelled as control, grey bars in panels B-H), or *Blos1*^*s*^ (blue bars in panels B-H) and either wt or mHTT-GFP under a doxycycline-inducible promoter. We induced expression using doxycycline (dox; 4.5 μM, 72 h), collected RNA, and measured *Xbp1* splicing by PCR followed by gel analysis. (C, D) Using samples from B, we measured the relative mRNA abundance of the RIDD targets *Hgsnat* and *Blos1* by real-time quantitative PCR (qPCR). (E, F) We induced expression of mHTT-GFP (4.5 μM dox, 72 h), then fixed and stained cells for LAMP1 to image LE/lysosomes (scale bar, 10 μm). F shows the quantification of 3 independent experiments (100-120 cells per treatment per experiment). (G) We induced expression of wt or mHTT-GFP (0.9 or 4.5 μM dox, 36 h), and measured relative *Blos1* abundance by qPCR. (H) We induced expression of mHTT as in G and quantified LE/lysosome localization as in E,F. For all graphs: independent experiments are indicated by symbol type, and bars show the average of 3-4 experiments. *, p < 0.05, paired t-test followed by Holm-Bonferroni correction for multiple comparisons.

Large aggregates of mHTT were visible in only 13% of control cells in these experiments (Fig. 2B), suggesting that extensive aggregation is not required to activate IRE1. Furthermore, when we used more mild induction conditions (0.9 μM doxycycline, 36 h) such that the mHTT-GFP fluorescence remained diffuse (Fig. 2A,B), cells still degraded *Blos1* mRNA and repositioned LE/lysosomes (Fig 1G,H). This is consistent with previous studies indicating that mHTT activates ER stress pathways by interfering with the retrotranslocation and proteasomal degradation of misfolded proteins from the ER (Duennwald and Lindquist, 2008; Leitman, Ulrich Hartl and Lederkremer, 2013), rather than through a mechanism dependent on extensive aggregation.

**Figure 2.**
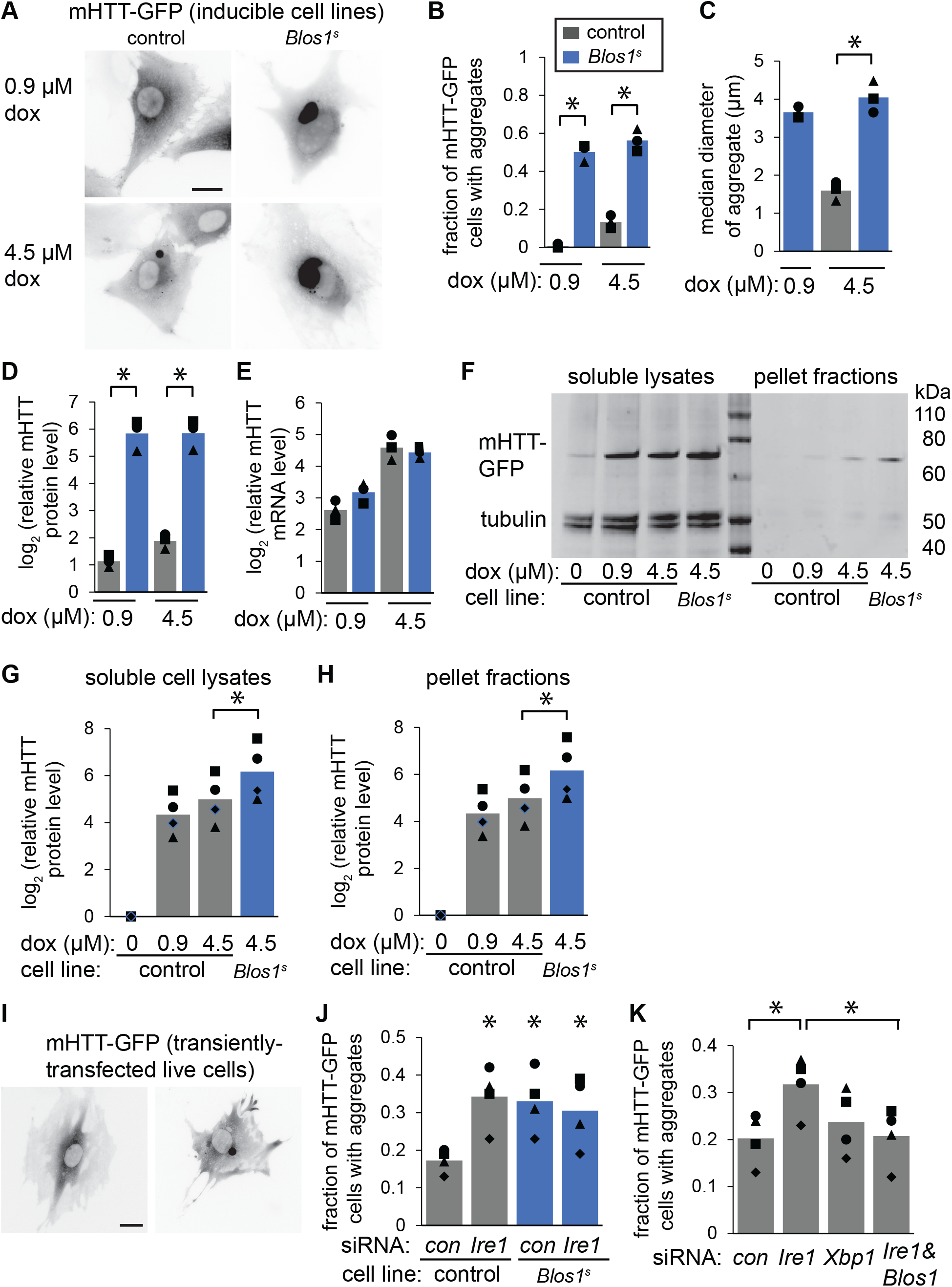
*Blos1* degradation prevents the accumulation of mHTT aggregates. (A-C) We induced expression (0.9 or 4.5 μM dox, 36 h) of mHTT-GFP in stable cell lines from Fig 1B-H. We then imaged live cells (representative images are in A, scale bar, 10 μm), quantified the fraction of cells with a prominent juxtanuclear aggregate (B) and measured the median diameter of the aggregate using ImageJ (C). (D-H) We induced expression of mHTT-GFP as in A-C and measured the median abundance of mHTT-GFP by flow cytometry (D), or collected RNA and measured relative mHTT-GFP mRNA abundance by qPCR (E), or measured the protein abundance by immunoblotting (F-H). (I-K) We used RNAi to deplete *Ire1, Xbp1*, and/or *Blos1* from MC3T3-E1 cells expressing *Rfp* (control, grey bars) or *Blos1*^*s*^ (blue bars) for 48 h, then transiently transfected mHTT-GFP, and imaged live cells 24 h post-transfection. Representative images in I show diffuse and aggregated mHTT-GFP (scale bar, 10 μm); we counted the fraction of fluorescent cells containing aggregates (J,K, 100 cells per replicate). For all graphs: independent experiments are indicated by symbol type, and bars show the average of 3 experiments. *, p < 0.05, paired t-test followed by Holm-Bonferroni correction for multiple comparisons.

Several lines of evidence suggest that RIDD of *Blos1* is important for degrading mHTT and avoiding the excessive accumulation of aggregates. Cells co-expressing *Blos1*^*s*^ were much more likely to contain juxtanuclear mHTT aggregates, and these aggregates were larger compared to those in control cells (Fig. 2A-C). To determine the extent of mHTT accumulation in these cells, we measured the fluorescence of its GFP tag by flow cytometry. *Blos1*^*s*^-expressing cells had a median mHTT-GFP signal over 25 times higher than control cells (Fig 2D), although the mRNA levels were similar between the two cell lines (Fig 2E). Immunoblot analysis confirmed the higher levels of mHTT in *Blos1*^*s*^-expressing cells (Fig. 2F-H), although the absolute abundance of mHTT is likely to be underestimated in cells containing aggregated proteins. We solubilized precipitate fractions from cell lysates by passing through a needle; again *Blos1*^*s*^-expressing cells contained higher levels of mHTT-GFP in the pellet fractions (Fig. 2F, H). Finally, depleting control cells of *Ire1*, but not *Xbp1*, by RNA interference (RNAi) increased the fraction of transiently-transfected mHTT-GFP-expressing cells containing aggregates, whereas mHTT aggregation in *Blos1*^*s*^-expressing cells was not affected by *Ire1* depletion (Fig. 2I-K). Co-depletion of both *Ire1* and *Blos1* restored the baseline level of mHTT aggregation (Fig. 2K), suggesting that the primary effect of IRE1 on mHTT aggregation is mediated by degradation of the *Blos1* mRNA.

Mutant HTT has previously been shown to be degraded by macroautophagy (MA). Consistently, treatment of our stable cell line that inducibly expresses mHTT-GFP with high levels of doxycycline (4.5 μM) led to higher levels of lipidated LC3B (LC3B-II, Fig. 3A-B), a marker for the induction of MA. However, at lower concentrations of doxycycline (0.9 μM), LC3B-II levels remained low and indistinguishable from cells without doxycycline. It was not possible to determine precisely whether the increase in LC3B-II levels in high doxycycline was due to increased induction of MA *vs*. less efficient degradation of LC3B-II following the fusion of autophagosomes with lysosomes, because inhibiting lysosomal function for the full timescale of the experiment led to extensive cell death. However, including chloroquine, which blocks the acidification of lysosomes, for the final 2 hours did lead to increased LC3B-II in all conditions (Fig. 3A-B). To test the idea that MA is responsible for degradation of mHTT only at high expression levels, we used RNAi to deplete the mRNA encoding the essential MA factor ATG7, induced mHTT expression with either 0.9 or 4.5 μM doxycycline, and measured the accumulation of mHTT-GFP by flow cytometry. In the high doxycycline conditions, *Atg7* knockdown led to a large increase in mHTT levels, whereas in the low doxycycline conditions, *Atg7* knockdown had no effect (Fig 3C). In contrast, knockdown of either *Hdac6* (which is important for trafficking aggregating proteins to the MTOC) or *Vps22* (which is important for degradation of ubiquitinated aggregates during ER stress (Bae *et al*., 2019)), led to accumulation of mHTT in both doxycycline concentrations (Fig 3C).

**Figure 3.**
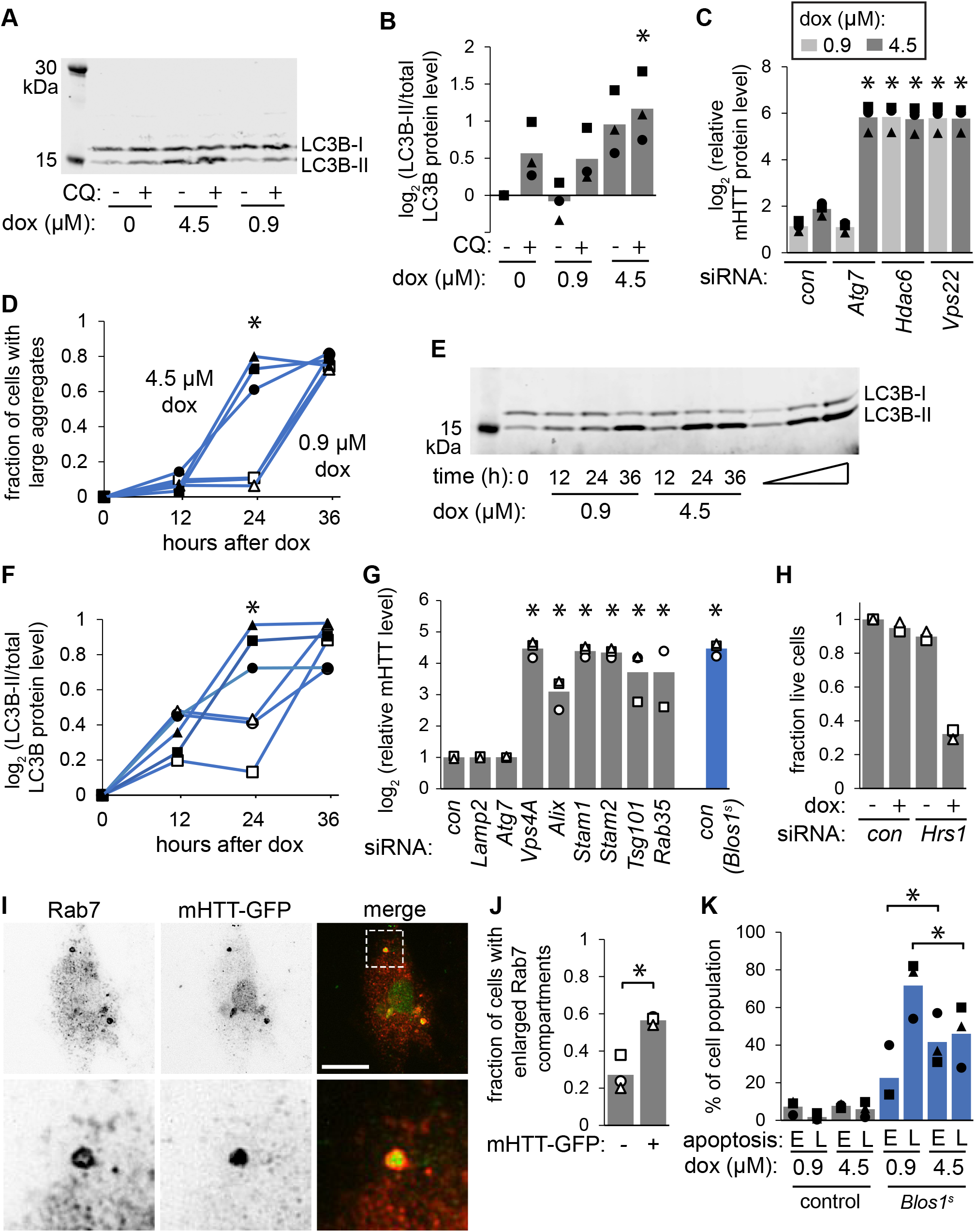
The degradation mechanism and toxicity of mHTT depends on expression level. (A-B) We induced expression of mHTT-GFP in control cells from Fig. 1B-H (0.9 or 4.5 μM dox, 36 h), with or without chloroquine (CQ, 60 μM) included for the final 2 h. We then measured LC3B processing by immunoblotting. Quantification of three independent immunoblots are shown in B. (C) We used RNAi to deplete MC3T3-E1 cells of the indicated factors, then induced mHTT-GFP expression (36 h) and measured the median fluorescence of cell populations by flow cytometry. *, p < 0.05 compared to the control knockdown at the same dox concentration. (D-F) We induced expression of mHTT-GFP in MC3T3-E1 cells expressing *Blos1*^*s*^ with either 0.9 μM (open symbols) or 4.5 μM (closed symbols) dox, and then measured the fraction of cells with large GFP aggregates (by microscopy, D) and the extent of MA induction (by LC3B immunoblot, E-F) over time. *, p < 0.05 comparing 0.9 vs 4.5 μM dox. The open triangle in E indicates increasing amounts of protein were loaded to test for signal intensity linearity. (G) We used RNAi to deplete MC3T3-E1 cells of the indicated factors, induced mHTT-GFP with low dox concentrations (0.9 μM, 36 h), and measured the median fluorescence as in B. For this panel, the data were analyzed by one-way ANOVA followed by Tukey’s Honest Significant Difference test (*, p < 0.05 compared to the control knockdown). (H) We used RNAi to deplete MC3T3-E1 cells of *Hrs1*, induced mHTT-GFP with low dox concentrations (0.9 μM, 36 h), aspirated floating cells from the plate and counted remaining living cells. (I-J) We transiently transfected MC3T3-E1 cells with mHTT-GFP (0.5 ug, 24 h), then treated briefly with digitonin to release soluble protein before fixing and staining for GFP and RAB7. Scale bar, 10 μm. We then counted the fraction of cells containing enlarged RAB7 structures in untrasfected vs GFP-containing cells. (K) We induced expression of mHTT-GFP in cells expressing *Rfp* (control, grey bars) or *Blos1*^*s*^ (blue bars) for 36 h, then measured the degree of apoptosis using annexin V and propidium iodide staining followed by flow cytometry. E=early apoptosis (staining with annexin V only), L=late apoptosis (staining with both annexinV and PPI). For all graphs: independent experiments are indicated by symbol type, and bars show the average of 3 experiments. *, p < 0.05, paired t-test followed by Holm-Bonferroni correction for multiple comparisons, except where noted. Controls confirming the depletion of correct targets by RNAi are in Fig S1.

These results suggest that mHTT induces MA and is degraded by MA only when expressed at high levels, potentially when larger aggregates begin to form. To further test this idea, we measured both aggregate formation (by microscopy) and MA induction (by immunoblot of LC3B-II) over time in cells expressing *Blos1*^*s*^, which accumulate aggregates when mHTT expression is induced with either low or high concentrations of doxycycline. The induction of MA coincided with the appearance of large, visible aggregates, after 24 hours of 4.5 μM doxycycline or 36 hours of 0.9 μM doxycycline (Fig 3D-F). We therefore propose that while MA is involved in mHTT degradation, a distinct pathway is necessary for its degradation at lower expression levels and/or smaller aggregation states, before large aggregates form. Furthermore, because *Blos1*^*s*^ affects mHTT degradation in both cases, RIDD of *Blos1* appears to enhance this MA-independent pathway.

To explore the mechanism of mHTT degradation in the low expression and sub-aggregation state, we depleted cells of various factors by RNAi, induced mHTT expression with 0.9 μM doxycycline, and measured its accumulation by flow cytometry. Similar to the depletion of *Atg7*, depletion of *Lamp2* (which is important for chaperone-mediated autophagy) did not affect mHTT levels (Fig 3G). However, we observed strong mHTT accumulation in cells depleted of factors involved in the endosomal sorting complex required for transport (ESCRT) pathway, which is responsible for the inward budding of vesicles into LE’s or multivesicular bodies (Henne, Buchkovich and Emr, 2011). The ESCRTs are composed of distinct complexes (ESCRTs −0, -I, - II, -III, and VPS4) that sort cargo and deform the limiting membrane of the endosome to allow for the formation of LEs, which then fuse with lysosomes for cargo degradation. In our experiments, mHTT accumulated following knockdown of the ESCRT-I factor *Tsg101*, the ESCRT-II factor *Vps22*, the ESCRT-III accessory factor *Alix*, and *Vps4A*, which is essential for the disassembly and recycling of ESCRT machinery (Fig 3C,G).

ESCRT-0 also appears to be critical for preventing the accumulation of mHTT. ESCRT-0 is a heterotetromer of two HRS and two STAM1 or STAM2 subunits, and it binds both endosomal membranes and ubiquitinated targets and initiates the multivesicular body pathway (Henne, Buchkovich and Emr, 2011). Depletion of *Stam1* or *2* led to dramatic accumulation of mHTT (Fig 3G), whereas depletion of the *Hrs1* led to cell death upon induction of mHTT expression (Fig 3H). Furthermore, depletion of *Rab35*, which encodes a small GTPase that regulates HRS (Sheehan *et al*., 2016; Vaz-Silva *et al*., 2018), also led to strong mHTT accumulation (Fig. 3G). Depletion of *Stam1*, but not *Atg7*, also led to the accumulation of mHTT aggregates 24 hours after transient transfection with low concentrations of the mHTT-GFP plasmid, confirming the effects measured by flow cytometry (Fig S1E).

Based on these data, we conclude that several core components of the ESCRT machinery are important for the degradation of mHTT. ESCRTs are involved in many cellular mechanisms, and directly or indirectly affect all three major autophagic pathways (Vietri, Radulovic and Stenmark, 2020), i.e. MA, chaperone-mediated autophagy, and endosomal microautophagy (eMI). However, the lack of dependence on MA or chaperone-mediated autophagy for mHTT degradation at low induction levels suggests that the ESCRTs contribute to mHTT degradation directly via eMI (Schuck, 2020). Although the mechanistic details are not yet well-understood for mammalian cells, eMI is characterized by the delivery of cytosolic cargo, such as glycolytic proteins, to LEs in a manner that relies on several ESCRT proteins, including TSG101 and VPS4 (Sahu *et al*., 2011; Vietri, Radulovic and Stenmark, 2020).

To further test a potential role for eMI in the degradation of mHTT-GFP, we examined its colocalization with RAB7, an LE marker, in cells transiently transfected under conditions where depletion of *Stam1* but not *Atg7* affects aggregate accumulation (0.5 μg of plasmid for 24 h, Fig. S1E). In these conditions, most cells display bright, diffuse GFP signal, obscuring any potential encapsulation by endocytic organelles. We therefore pre-permeabolized cells with digitonin (60 μg/mL, 4C, 15 min) before fixing and staining for GFP and RAB7. This approach reduced the GFP signal and revealed several detergent-resistant GFP foci in most cells, whereas RAB7 staining appeared similar to that in cells without pre-permeabolization. Cells transfected with mHTT-GFP were more likely than untransfected cells to contain enlarged or more prominent/brighter RAB7 foci (Fig 3I,J), usually 1-2 per cell. Strikingly, these RAB7 structures co-localized with mHTT-GFP in 71 % of the cells (-/+ 0.08%, in an average of 3 independent experiments with 93-96 cells per replicate) (Fig 3I). These results are consistent with the idea that mHTT-GFP is encapsulated in LEs during its degradation prior to forming large, juxtanuclear aggregates. As expected due to the transient nature of cargo-containing LEs, these did not account for most of the detergent-resistant GFP foci, which may represent aggregates or other membrane-bound structures, although we did not detect any colocalization with LC3 under these conditions.

We next carried out apoptosis assays to test the relative toxicity of mHTT in both control and *Blos1*^*s*^-expressing cells after treatment with low or high doxycycline concentrations. Control cells were much more resistant to apoptosis, indicating that degrading *Blos1* mRNA not only prevents the accumulation of mHTT aggregates but also protects cells from death. Surprisingly, *Blos1*^*s*^-expressing cells treated with 0.9 μM dox were positive for both annexin V and propidium iodide, indicative of late apoptosis, whereas cells treated with 4.5 μM dox were more likely to be stained for annexin V only, indicative of early apoptosis (Fig. 3K). These differences cannot be explained by the amount of mHTT or the number of aggregates at the time of the assay (36 h after doxycycline addition), as these were indistinguishable between the two doxycycline treatments (Fig 2D and 3D). However, due to the delay in forming large aggregates in the lower expression regime (Fig 3D), we suggest that these cells are exposed to a more toxic, sub-aggregation form of mHTT for a longer time compared to the high expression regime where mHTT accumulates in large aggregates more rapidly.

To test the generality of these findings and the ability of *Blos1* regulation to enhance mHTT degradation in a neuronal cell line, we transfected wt or mHTT-GFP into mouse Neuro2A (N2A) cells, with or without *Blos1*^*s*^. As in the MC3T3-E1 cells, expression of mHTT led to the degradation of the *Blos1* mRNA, and co-expression of *Blos1*^*s*^ led to increased accumulation of mHTT but not wtHTT (Fig 4A-B). We then asked whether mHTT is degraded by eMI and/or MA in these cells. In control cells not expressing *Blos1*^*s*^, knockdown of the ESCRT-0 factor *Stam1*, but not the MA factor *Atg7*, led to increased mHTT accumulation (Fig 4C). There was no difference between mHTT levels in control vs. *Blos1*^*s*^ cells after depletion of *Stam1*. In contrast, *Atg7* knockdown led to increased mHTT accumulation beyond the effects of *Blos1*^*s*^ expression. Taken together, these data suggest that the effect of *Blos1* regulation is primarily on eMI. When this pathway is crippled by *Blos1*^*s*^ expression, knockdown of *Stam1* has no further effect, whereas knockdown of *Atg7* (which normally does not affect mHTT degradation in these cells) exacerbates *Blos1*^*s*^ expression by preventing mHTT degradation by MA.

**Figure 4.**
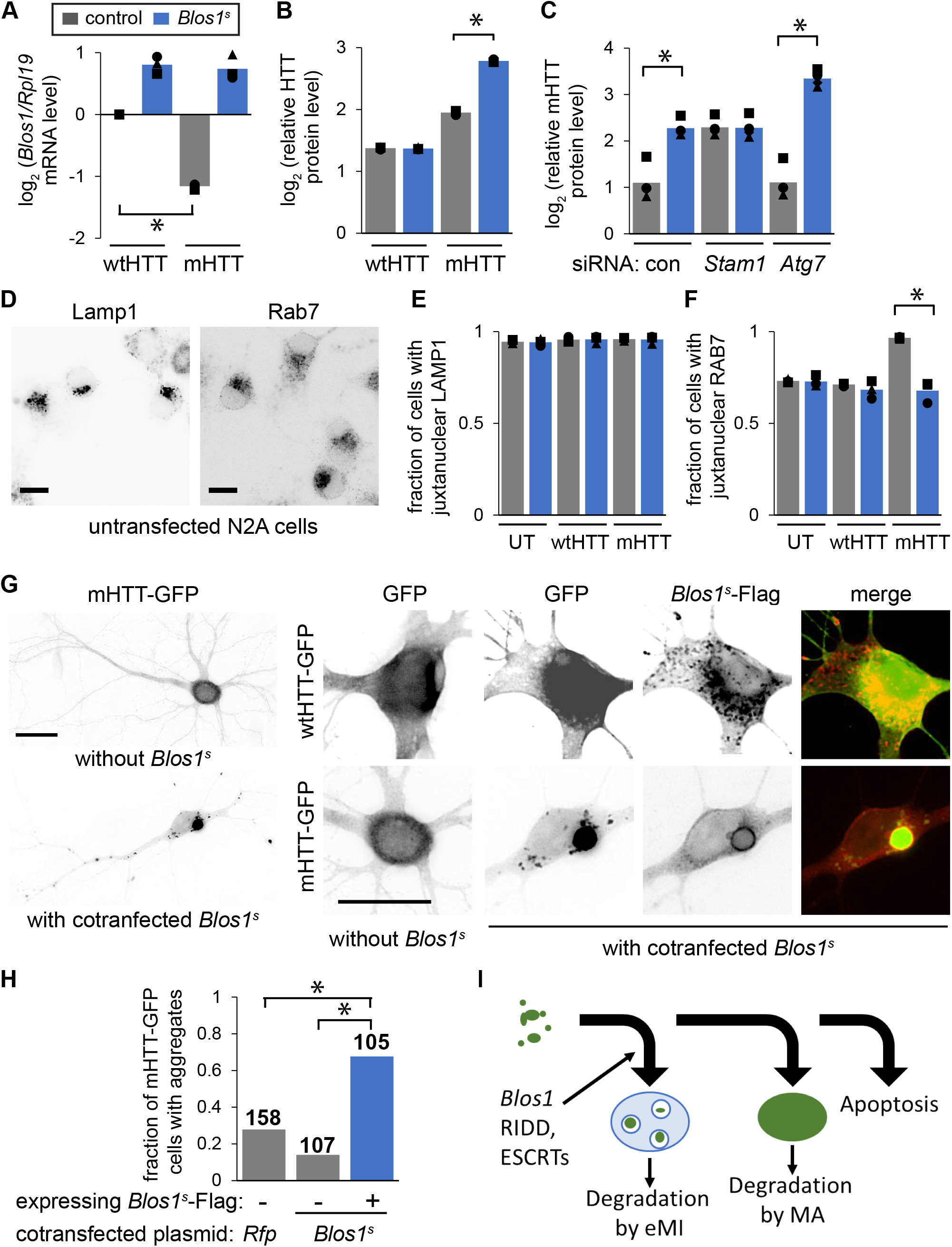
*Blos1* regulation protects cells from mHTT accumulation in N2A cells and primary rat hippocampal neurons. (A,B) We cotransfected N2A cells with *Rfp* (labelled as control, grey bars), or *Blos1*^*s*^ (blue bars) and either wt or mHTT-GFP under a constitutive promoter. After 48 h, we measured *Blos1* mRNA levels by qPCR (A) and the accumulation of HTT-GFP by flow cytometry (B). (C) We used RNAi to deplete N2A cells of *Stam1* or *Atg7*, then transfected and measured the accumulation of HTT-GFP as in B. (D-F) We compared untransfected N2A cells to those transfected as in A-B. We stained cells with antibodies for LAMP1 (D,E) or RAB7 (D,F) and scored the fraction of cells with juxtanuclear foci. Scale bars, 10 μm. (G,H) We isolated primary rat hippocampal neurons from E18 embryos, transfected with plasmids encoding wt or mHTT-GFP and either *Rfp* (as a control) or *Blos1*^*s*^*-flag*, fixed and stained with antibodies for GFP and FLAG (to identify co-transfected cells). Representative images are shown in G (scale bars, 20 μm.) In H, we scored the indicated number of neurons (4 transfections per condition) for the presence of large HTT-GFP aggregates. *, p < 0.05, Fisher’s exact test. Similar results were seen in independent neuron cultures. (I) Model for the degradation of small mHTT aggregates. For all graphs: independent experiments are indicated by symbol type, and bars show the average of 3 experiments. *, p < 0.05, paired t-test followed by Holm-Bonferroni correction for multiple comparisons (except where noted).

Unlike in MC3T3-E1 cells, LAMP1 structures in the N2A cells were predominantly juxtanuclear even in untransfected cells or cells transfected with wtHTT, and this localization was unaffected by *Blos1*^*s*^ expression (Fig. 4D, E). RAB7 structures, in contrast, were peripherally distributed in a higher fraction of cells, and did appear to be more juxtanuclear upon expression of mHTT but not wtHTT (Fig 4D, E). *Blos1*^*s*^ expression prevented this repositioning, similar to our results in MC3T3-E1 cells. Because LAMP1 stains both lysosomes and LEs, whereas RAB7 is typically more specific to LEs, the repositioning of LEs may account for the effects of *Blos1* degradation, at least in the case where lysosomes are already available near the MTOC. This is consistent with a primary effect of *Blos1* on eMI. Alternatively, as suggested by the large fraction (>50%) of untransfected cells displaying juxtanuclear RAB7 structures, *Blos1* regulation may affect mHTT accumulation in ways unrelated to the simple, global trafficking and localization of these organelles. In support of this idea, BORC, the BLOS1-containing complex responsible for LE/lysosome trafficking, has also been shown to regulate autophagosome-lysosome fusion (Jia *et al*., 2017) and LE/lysosome size (Yordanov *et al*., 2019). Furthermore, BLOS1 itself participates in a variety of related cellular functions as outlined above, both through its role in the biogenesis of lysosome related organelles complex (BLOC1) and potentially independently (Scott *et al*., 2018).

Finally, to test whether *Blos1* degradation reduces the accumulation of mHTT in primary neurons, we isolated hippocampal neurons from rat E18 embryos and transfected at 15 days in vitro (DIV) with either wt or mHTT-GFP, along with either *Blos1*^s^-Flag or RFP as a control. We then fixed and stained the neurons at DIV 19 using antibodies for GFP and FLAG (Fig. 4G, H). Neurons expressing wtHTT-GFP displayed diffuse, bright GFP signal, whereas neurons expressing mHTT-GFP contained either diffuse signal or large juxtanuclear aggregates. Neurons co-expressing *Blos1*^*s*^ (identified by FLAG staining) were more likely to contain large aggregates, compared with neurons on the same coverslip lacking FLAG signal, or neurons cotransfected with the control RFP (Fig. 4G, H). These data suggest conserved mechanisms of aggregate clearance in cultured cells and neurons.

Our overall model is that prior to forming large, juxtanuclear aggregates, mHTT is degraded by eMI, in a manner that depends on the degradation of *Blos1* by the RIDD pathway, potentially due to the repositioning of LEs toward the MTOC (Fig. 4F). When this degradation pathway is compromised (as in cells expressing *Blos1*^*s*^), mHTT accumulates in juxtanuclear aggregates and MA is induced. Although the regulation of *Blos1* has not been explored in neurodegenerative diseases, the aggregation-prone tau protein, which is involved in the pathogenesis of Alzheimer’s disease, has also been shown to be degraded by both eMI and MA (Caballero *et al*., 2018; Vaz-Silva *et al*., 2018). More generally, dysfunction of the ESCRT machinery, including ESCRT-0, has been linked to neurodegeneration and the accumulation of intracellular protein aggregates (Oshima *et al*., 2016; Kaul and Chakrabarti, 2018).

Why are different degradation pathways used for the clearance of mHTT? It is likely that MA is necessary for the removal of aggregates once they reach a certain size, but why cells would use eMI rather than MA in the case of monomers or smaller oligomers is an open question. We speculate that by enclosing these smaller aggregates within LEs, neurons retain the option of not only degrading these toxic species within their own lysosomes, but also secreting them outside the cell within exosomes, upon fusion of LEs with the plasma membrane. This mechanism, which is emerging as a common feature of neurodegenerative diseases, may help to protect neurons from toxicity when lysosomal capacity is not sufficient, while also potentially explaining the propagation of aggregates to target cells that take up these vesicles (Hill, 2019; Kalluri and LeBleu, 2020).

## Acknowledgments

We thank Jason Perry for technical assistance, and Markus Babst and members of the Hollien lab for discussions. This research was supported by NIH R35 GM119540. The authors declare no competing financial interests.

## Author contributions

Conceptualization, investigation, review & editing: all authors. Original draft: DB and JH.

## Figure Legends

**Figure S1.**
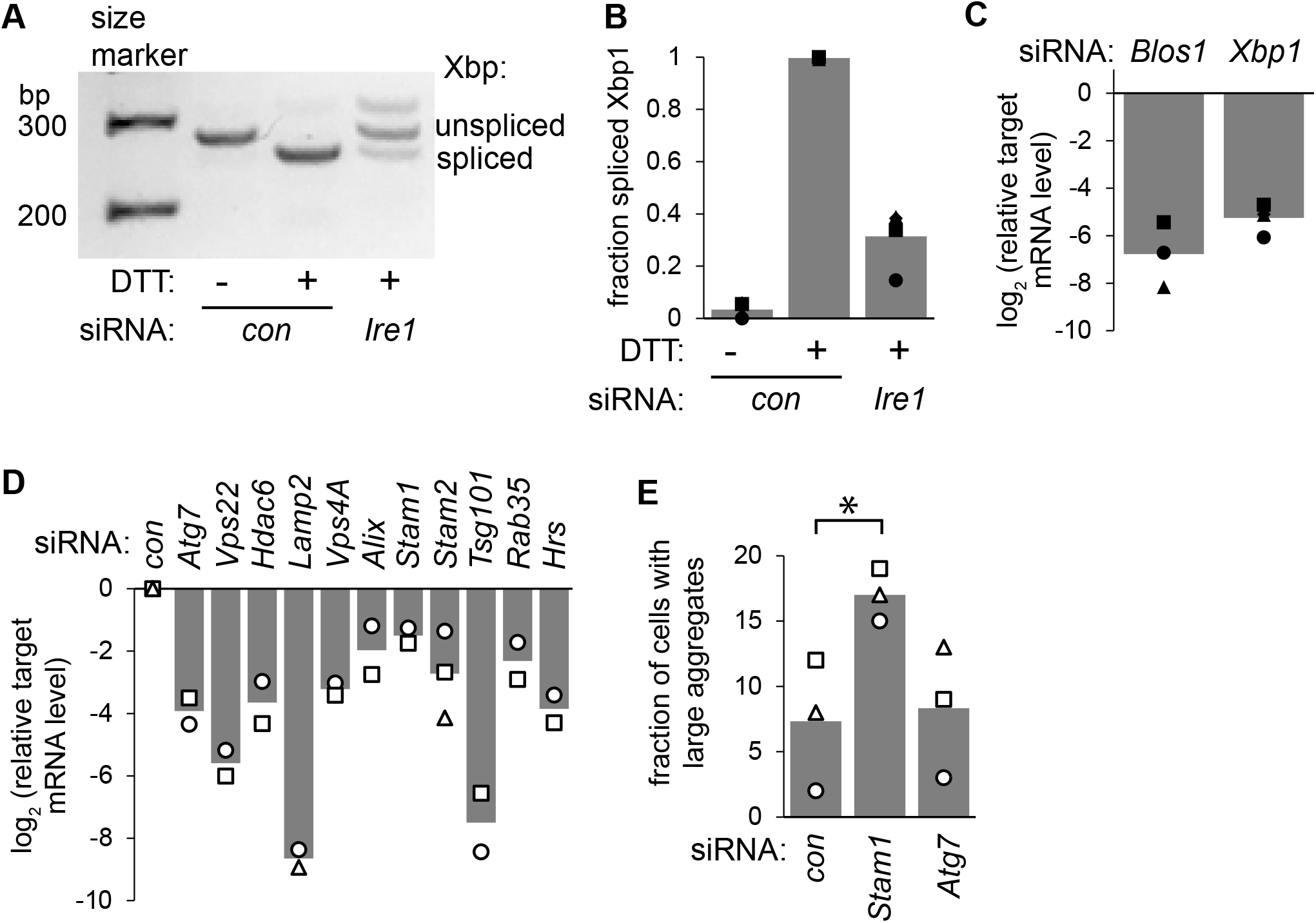
RNAi controls and confirmation experiments. (A-C) In parallel to the RNAi experiments in Fig. 2J,K, we treated a subset of cells with DTT (2 mM, 2 hrs) to assess the effects of *Ire1* depletion on its *Xbp1* splicing activity, and collected RNA from all samples to measure the knockdown efficiency. For *Ire1*-depleted cells, we measured the splicing of *Xbp1* mRNA by PCR followed by gel analysis (a representative gel is shown in A, and quantified splicing efficiency is shown in B). For *Blos1* and *Xbp1* depletion (C), we measured levels of these target mRNAs by qPCR, divided by the levels of the housekeeping control mRNA *Rpl19*, and divided target/ *Rpl19* levels in the knockdown samples by that in cells treated with control siRNAs. (D) RNAi efficiency for depletion experiments from Fig. 3C,G was measured by qPCR, as in S1C. (E) We used RNAi to deplete *Stam1* or *Atg7* from MC3T3-E1 cells for 48 h, then transiently transfected mHTT-GFP (0.5 μg plasmid), and counted the fraction of GFP-expressing cells that contained aggregates after 24 h (100 cells per replicate, *, p < 0.05, paired t-test).

## Materials and Methods

### Cell Culture, plasmids, and transfections

We cultured MC3T3-E1 cells in MEMα with nucleosides, L-glutamine,and no ascorbic acid (Life Technologies) supplemented with 10% FBS at 37 °C and 5% CO2. We cultured N2A cells in DMEM supplemented with 10% FBS. Both cell lines were purchased from American Type Culture Collection (ATCC).

We subcloned PCR products encoding exon 1 of either wild-type (23 CAG repeats) or mutant (145 CAG repeats) Huntingtin (HD Community Biorepository, ref# CHDI-90000038 for wild-type Huntingtin and CHDI-90000040 for mutant huntingtin), followed by GFP, downstream of the human EF1α promoter (for constitutive expression in transiently-traasnfected cells) or the doxycycline-inducible tight TRE promotor (for inducible expression in stably-transfected cells). The *Blos1*^*s*^ construct described previously (Bae *et al*., 2019) contains a silent point mutation (G360C) in the coding sequence that prevents its degradation by RIDD (Moore and Hollien, 2015).

To generate the MC3T3-E1 stable cell lines used in this study, we used control cells (expressing *Rfp*) and *Blos1*^*s*^-3x*Flag*-expressing cells described previously (Bae *et al*., 2019), and carried out additional transfections using Lipofectamine 2000 (Invitrogen). Into each cell line, we transfected a TetR plasmid (addgene: pMA2640 (Alexeyev *et al*., 2010)), which expresses the reverse tetracycline-regulated transactivator to allow for doxycycline-inducible expression, and selected for stable expression using blasticidin (5 μg/mL). We then transfected the wt or mHTT-GFP plasmid and selected using puromycin (2 μg/mL). We maintained cells in these antibiotics until one passage before each experiment.

We transiently transfected MC3T3 or N2A cells using Lipofectamine 2000 or 3000 (Invitrogen). For each experiment we transfected plasmids expressing with wt or mHTT-GFP under the constitutive EF1α promoter, along with plasmids expressing either *Rfp* (as a control) or *Blos1*^*s*^*-3xFlag* where indicated. We allowed the cells to recover for 24 hours (Fig 3 and S1) or 48 hours (Fig 2, 4) before assaying.

### RNA interference

For siRNA-mediated silencingexperiments in MC3T3-E1 cells, we used RNAiMAX (Invitrogen) to transfect two siRNAs (Sigma-Aldrich) for each target mRNA. As a control we transfected siRNAs designed to not target any mammalian mRNAs (Qiagen). For the double-knockdown of *Ire1* and *Blos1* in Fig 2K, we included the standard 450 ng of *Ire1* siRNAs plus 90 ng of *Blos1* siRNAs, which we had previously determined are effective in depleting *Blos1* mRNA at lower transfection amounts (confirmed in Fig. S1C). Forty-eight hours after siRNA transfection, we transiently transfected the constitutive HTT-GFP plasmids or induced expression in the stable cell lines using doxycycline.. RNAi in N2A cells was carried out in the same way, except that we allowed cells to recover for 24 h after siRNA transfection before transfecting with plasmids, followed by 48 h recovery before assaying.

### Primary hippocampal neuron culture and transfections

We isolated primary rat hippocampal neurons from E18 embryos from pregnant Sprague Dawley rats (Charles River Laboratory) as previously described (Shepherd *et al*., 2006). Dissected hippocampi were dissociated using DNase (0.01%, Sigma-Aldrich) and papain (0.067%, Worthington Biochemicals), triturated with glass polished pipettes, and strained through 70-μm pore-size cell strainer (Thermo Fisher Scientific) to obtain single-cell suspension. Neurons were plated on poly-L-lysine (0.2 mg/mL; Sigma-Aldrich) coated glass coverslips (Carolina Biological Supply) at 60,000 cells/mL in 12-well plates (Greiner Bio-One). Neurons were initially plated in neurobasal media (Gibco) containing 5% fetal bovine serum (Gibco), 1% GlutaMAX (Gibco), 1% penicillin/streptomycin (Gibco), and 2% SM1 supplement (Stem Cell Technologies), and subsequently fed with astrocyte-conditioned BrainPhys media (Stem Cell Technologies) and 2% SM1 every three days thereafter. Neurons were maintained in an incubator at 37°C and with 5% CO2.

We transfected neurons at 15 days in vitro using lipofectamine 2000 (Thermo Fisher). For one well, 1.5 ug of each plasmid was complexed with lipofectamine 2000 as per the manufacturer’s protocol. Neurons were transfected for 1 hour in 0.2 um filtered Minimum Essential Media (Thermo Fisher) supplemented with 2% GlutaMax (Gibco), 2% SM1 (Stem Cell Technologies), 15mM HEPES (Gibco), 1mM sodium pyruvate (Gibco), and 33 mM glucose (pH=7.2) at 37°C without CO2. After transfection, neurons were given 96 hours in growth media at 37°C/ 5% CO2 to allow sufficient recovery and expression of the HTT proteins. Neurons were then washed with 1x phosphate-buffered saline (PBS) once and fixed using 4% formaldehyde (Thermo Fisher)/4% sucrose (VWR) in 1x PBS for 15 minutes at room temperature. Permabolization and staining were carried out as described below for cultured cells.

### Immunostaining and microscopy

For immunostaining, we grew cells on glass coverslips, fixed using 4% paraformaldehyde and 1 mM MgCl in PBS (37 C, 15 min), and permeabilized using 0.2% Triton X-100 and 1 mM MgCl in PBS (room temperature, 20 min). We then incubated cover slips in blocking solution (2% BSA, 0.02% Tween-20, and 1 mM MgCl in PBS, 10 min), then with primary antibodies in blocking solution (room temperature, 1 hour). We used primary antibodies for GFP (Aves labs GFP1010, 1:1000), alpha-TUBULIN (Cell Signaling 2144, 1:25), LAMP (DSHB, 1D4B-s, 2 μg/mL), RAB7 (Cell Signaling 9367, 1:100), or FLAG (Sigma F3165, 1:500). We washed 3 times (0.02% Tween-20, 1 mM MgCl, PBS), incubated with secondary antibodies (1 hour), and washed again. We then mounted the coverslips on slides in ProLong Diamond Antifade mountant with DAPI (Invitrogen). For live cell imaging, we plated cells in glass-bottom dishes.

To image cells, we used an Olympus IX-51 inverted microscope with a 60x (NA 1.25) oil objective at room temperature and either a Q-imaging Qicam (SN Q25830; acquisition sotware QCapturePro 6.0) or an Olympus DP23 monochrome camera (acquisition software CellSens Standard v3). To quantify LE/lysosome positioning, we assigned random file names and had a researcher blinded to the conditions score each cell. Cells with >50% of the LAMP1 or RAB7 foci located next to and on one side of the nucleus were counted as displaying predominantly juxtanuclear LE/lysosomes. We scored approximately 100 cells for each condition in each experiment and repeated each experiment at least 3 times.

### Quantitative real-time RT-PCR and *Xbp1* splicing assays

We collected cells and isolated total RNA using Quick RNA Miniprep kits (Zymo Research). We synthesized cDNA with 700 ng-2 μg total RNA as a template, a T18 primer, and Moloney murine leukemia virus reverse transcriptase (New England Biolabs). Using a QuantStudio 3 real-time quantitative PCR machine (Life Technologies), we measured relative amounts of specific mRNAs with SYBR green as the fluorescent dye. All measurements were done in triplicate, and we quantified by comparing to serially diluted standard curve samples. Target mRNA primers are listed in Table S1. We divided the abundance of each target mRNA by that for ribosomal protein 19 (*Rpl19*) mRNA from the same sample. For *Xbp1* splicing, we amplified cDNA with primers surrounding the regulated *Xbp1* splice site (AGAAGAGAACCACAAACTCCAG and GGGTCCAACTTGTCCAGAATGC) and ran the PCR products on a 2% agarose gel. We quantified the relative intensities of the spliced and unspliced *Xbp1* bands using ImageJ.

### Immunoblotting

We trypsinized and collected cells, and lysed in RIPA buffer (25 mM Tris, pH 7.6, 150 mM NaCl, 1% NP-40, 1% Na-deoxycholate, and 0.1% SDS) with protease and phosphatase inhibitors (Thermo Fisher Scientific). We removed insoluble material by centrifugation, and for GFP immunoblots we resuspended the pellets in equivalent volumes of SDS sample buffer and passed through a needle 10 times to solubilize. We then resolved proteins using either 4-12% (for GFP) or 12% (for LC3) polyacrylamide NuPage Bis-Tris gels. We transferred the proteins to nitrocellulose and incubated for 1 hour in blocking buffer (5% BSA, 0.05% Tween20, 0.01% Triton X-100, and TBS) at room temperature. We then probed using antibodies for GFP (Invitrogen Molecular Probes A6455, 1:7500), TUBULIN (Cell Signaling 2144, 1:5000), or LC3B (Sigma-Aldrich L7543, 1:1000) at 4 C overnight, washed, and incubated with a secondary antibody (LiCor 926-32211, 1:10,000, 1 hour, RT). We used a LiCor Odyssey CLx Imager to scan the blot and quantified band intensities using the LiCor Image Studio software. For GFP immunoblots we divided the GFP signal by the tubulin signal in soluble samples. For LC3 we divided LC3B-II (processed, lipidated LC3B) band intensities by the sum of the intensities for unprocessed LC3B-I and processed LC3B-II.

### Flow cytometry and Annexin V/PI apoptosis assays

For measuring HTT-GFP abundance, we used a CytoFLEX 5 flow cytometer (Beckman Coulter) and the CytExpert version 2.3 software. For each sample in each experiment, we measured the median GFP intensity in a minimum of 4800 live cells and divided by the signal for untransfected cells.

To measure apoptosis, we used the Annexin V-Alexa Fluor 684 apoptosis assay kit (Invitrogen) according to the manufacturer’s protocol, except that we used 5-fold less than the suggested annexin concentration to avoid excessive staining of control cells. We trypsinized and collected cells by centrifugation (800 xg, 5 min, RT), incubated in Annexin V (1:100, 15 min, 37 C) and Propidium Iodide (1 ug/mL, 2 min, 37 C), and measured fluorescence intensities by flow cytometry as above.

**Table S1.**
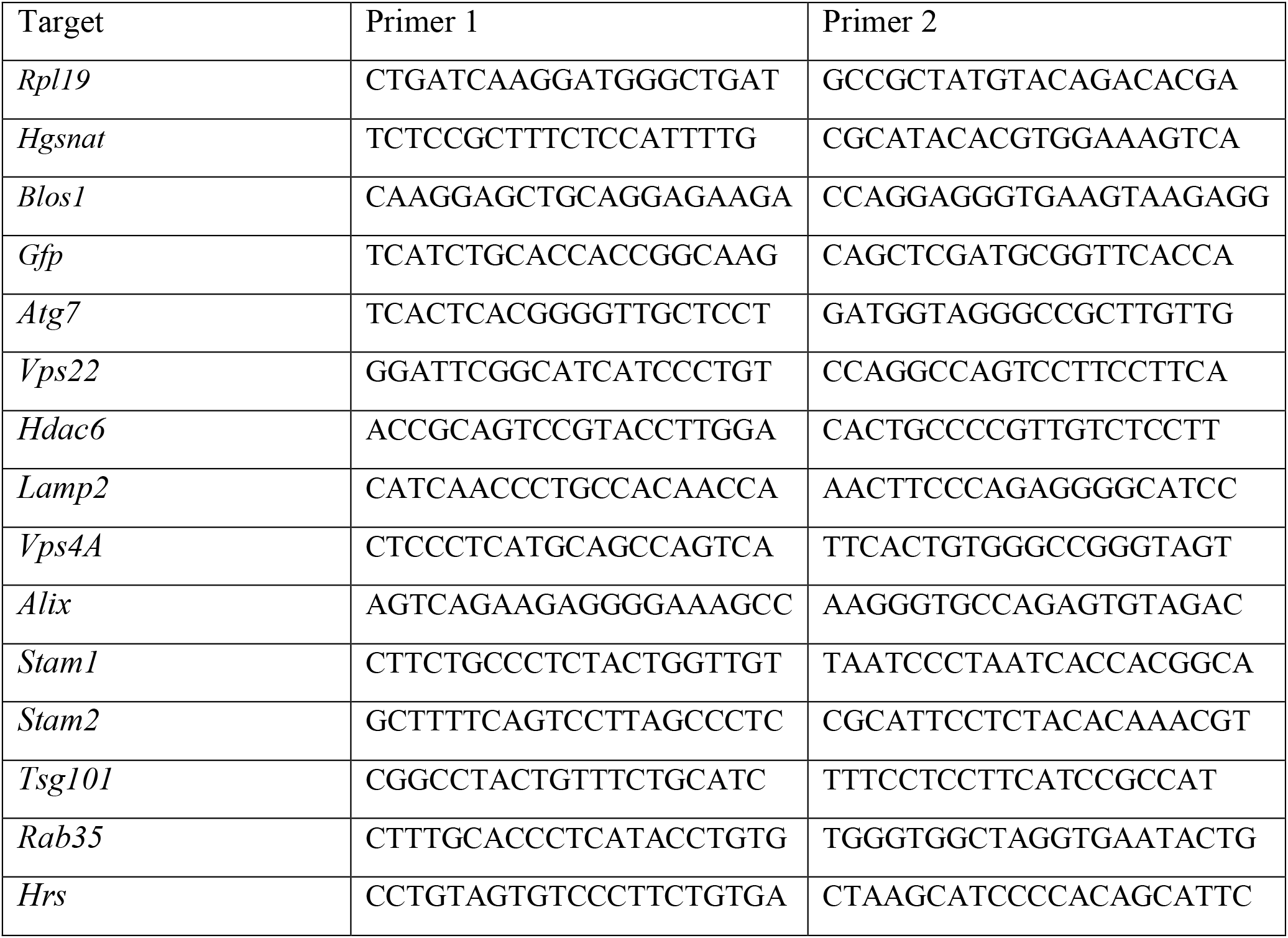
Primers used for quantitative PCR.

